# Imaging dendritic spines in the hippocampus of a living mouse by 3D-STED microscopy

**DOI:** 10.1101/2023.02.01.526326

**Authors:** Stéphane Bancelin, Luc Mercier, Johannes Roos, Mohamed Belkadi, Thomas Pfeiffer, Sun Kwang Kim, U. Valentin Nägerl

## Abstract

STED microscopy has been used to address a wide range of neurobiological questions in optically well-accessible samples like cell culture or brain slices. However, the application of STED to deeply embedded structures in the brain of living animals remains technically challenging. In previous work, we established chronic STED imaging in the hippocampus *in vivo* but the gain in spatial resolution was restricted to the lateral plane. In this study, we report on extending the gain in STED resolution into the optical axis to visualize dendritic spines in the hippocampus *in vivo*. The approach is based on a spatial light modulator to shape the focal STED light intensity in all three dimensions and a conically shaped window that is compatible with an objective that has a long working distance and a high numerical aperture. Moreover, we corrected distortions of the laser wavefront to optimize the shape of the bottle beam of the STED laser, which is required for 3D-STED microscopy. In summary, we present a methodology to improve the axial resolution for STED microscopy in the deeply embedded hippocampus *in vivo*, facilitating longitudinal studies of neuroanatomical plasticity at the nanoscale in a wide range of (patho-)physiological contexts.

## Introduction

The hippocampus is a deeply embedded brain region, which plays a critical role in encoding new memories in the mammalian brain. In the hippocampus, as elsewhere in the mammalian brain, pyramidal neurons receive most of their excitatory synaptic input at dendritic spines, which are small protrusions in the postsynaptic membrane that house the postsynaptic signaling machinery including glutamate receptors. Structural and functional plasticity of dendritic spines is a fundamental neurobiological process that underlies all higher brain functions like memory, thought and action (1, 2), while spine dysfunction is closely associated with neuropsychiatric and neurodegenerative disorders, such as autism and Alzheimer’s disease (3).

2-photon fluorescence microscopy provides high depth penetration and optical sectioning in turbid media (4), making it the standard technique for imaging in acute brain slices (5, 6) and intact brains (7–9). Over the last 20 years, it has transformed our understanding of the structure and function of dendritic spines in the mouse model of the brain (10).

Up to now, most *in vivo* studies of dendritic spines have been limited to superficial layers of the cortex (somatosensory, motor, visual cortex) (11–13), mainly because of the challenge to optically reach more deeply embedded structures. For example, the hippocampus is more than 1 mm below the cortical surface of the mouse brain. Only a few groups have ventured into imaging hippocampal spines *in vivo*, relying either on surgical resection of the overlaying cortex (14, 15) or a micro-endoscope with a gradient-index lens (16).

However, 2-photon microscopy is a diffraction-limited approach, which offers at best a spatial resolution of around 350 nm laterally and 1 *μm* axially, falling substantially short of visualizing several key neuro-anatomical structures and spaces, including spine necks, axon shafts, astroglial processes and synaptic clefts, whose sizes can range well below 100 nm. As a consequence, there is a great need to develop and improve super-resolution imaging techniques that could be applied to deeply embedded regions like the hippocampus. Among the various super-resolution methods, Stimulated Emission Depletion (STED) microscopy (17) is currently the only one that has been successfully applied in vivo (18–21), notably in the hippocampus (22).

STED microscopy is based on laser-scanning microscopy where a Gaussian laser beam is focused to a small spot generating the fluorescence signal used to construct the image. In addition, there is a second laser (the STED laser), which exerts the opposite effect, namely it de-excites molecules by the process of stimulated emission. By spatially shaping the focal STED intensity like a donut (17), it is possible to suppress the spontaneous fluorescence in the peripheral region of the focal spot, narrowing down the effective point spread function (PSF) in the x-y plane by up to an order of magnitude (23). By delivering STED light also above and below the focal region (a profile referred to as ‘bottle beam’), it becomes possible to constrict the fluorescence along the optical axis as well (24, 25).

In STED microscopy, spatial resolution and signal-to-noise ratio (SNR) of the images crucially depend on the quality of the PSF of the STED beam (26). Yet, maintaining a high-quality PSF inside light scattering brain tissue poses several challenges. This problem is particular evident in the context of in vivo imaging, where a variety of biological and mechanical constrains stand in the way of ensuring sufficiently good optical conditions for STED imaging. Notably, long working distance water immersion objectives typically used for *in vivo* imaging are not optimal for STED, where objectives with high numerical aperture (NA) and oil or glycerin as immersion medium are preferred. In addition, the mechanical stability of the cranial window attachment as well as of the brain itself are crucial, since any vibrations stemming from muscle contractions, pulmonary breathing and blood pulsations can produce motion artefacts and diminish image quality.

The bottle beam PSF used for the axial gain in resolution is particularly sensitive to optical aberrations and misconfigurations in the beam path. Notably, the modified cranial window used to image the hippocampus (15, 23) reduced the effective NA, which prevented the use of a bottle beam profile for 3D-STED in our previous study based on a cylindrically shaped ‘hippocampal window’ (22). In this work, we propose the use of a conically shaped window, specifically designed to maintain the bottle beam profile, while minimizing the size of the surgical resection of the overlaying cortex.

## Material and Methods

### 2P-STED microscope

Imaging was performed using a custom-built upright laser-scanning fluorescence microscope, as previously described (22, 26, 27). In brief, the 2-photon excitation beam (100 fs, 80 MHz, 900 nm) was provided by a femtosecond Titanium:Sapphire laser (Tsunami, Spectra-Physics) pumped by a high power cw DPSS laser (Millennia EV 15, Spectra-Physics) sent through a Pockels cell (302 RM, Conoptics) to control the excitation power. The STED beam (592 nm, 700 ps, 80 MHz) was provided by another pulsed laser (Katana 06 HP, NKT Photonics) whose power was adjustable using a half-wave plate and a polarization beam splitter. Both lasers were synchronized and temporally overlapped using commercial electronics (‘Lock-to-clock’, Model 3930 and 3931, Spectra-Physics).

The STED beam was passed via a spatial light modulator (SLM) (3D module, Abberior Instruments) to modulate the wavefront in a way and achieve a donut or bottle beam intensity distribution of the STED light in the focal plane. Half and quarter-wave plates (*λ*/2 and *λ*/4) were used to produce a left-handed circular polarization at the entrance pupil of the objective. Both 2-photon excitation and STED beams were combined using a long-pass dichroic mirror (DCSPXRUV – T700, AHF). Appropriate lens combinations were used to conjugate the SLM on a telecentric scanner (Yanus IV, TILL Photonics), which was then imaged on the back focal plane of the objective (CFI Apo NIR 60x W, NA 1.0, Nikon) mounted on a z-focusing piezo actuator (Pifoc 725.2CD, Physik Instrumente). This objective provided the necessary working distance to bridge the physical distance between the surface of the brain and the deeply embedded hippocampus, while still offering a relatively high numerical aperture conducive for high-resolution imaging.

The epi-fluorescence signal was de-scanned, separated from the incident beams using a long-pass dichroic mirror (580 DCXRUV, AHF) and detected by an avalanche photodiode (SPCM-AQRH-14-FC, Excelitas) with appropriate notch (594S-25, Semrock) and bandpass filters (680SP-25, 520-50, Semrock) along the emission path. Signal detection and hardware control were performed via a data acquisition card (PCIe-6259, National Instruments) and the Imspector software (Abberior Instruments).

To assess donut or bottle beam quality, a pellicle beam splitter (BP145B1, Thorlabs) was flipped into the beam path to detect the signal reflected by gold beads (150 nm Gold nanospheres, Sigma Aldrich) using a photomultiplier tube (MD963, Excelitas). In the following, 2D-STED, z-STED and 3D-STED will refer to images acquired using only a donut, only a bottle beam or a combination of the two beams. Optical resolution was assessed by imaging fluorescent beads (yellow-green fluorescent beads, 40 nm or 170 nm in diameter, Invitrogen) immobilized on glass slides.

### Animal experimentation

We used adult female and male transgenic mice (*Thy*1–*H^tg/+^*, 3 – 12 months old) where a subset of hippocampal neurons was fluorescently labeled with YFP (28). Heterozygous mice were used with sparse yet robust cytosolic labelling. The mice were group-housed by gender at a 12/12h light/dark cycles. All procedures were in accordance with the Directive 2010/63/EU of the European Parliament and approved by the Ethics Committee of Bordeaux under agreement number #8899.

### Hippocampal window implantation

Chronic hippocampal windows were implanted as described previously (14, 15, 22, 29) to provide optical access to the *stratum oriens* and *stratum pyramidale* of the CA1 region of the hippocampus. In brief, mice were anesthetized with isoflurane (2%) and received intraperitoneal injections of analgesic (buprenorphine, 0.05 mg/kg) and anti-inflammatory drugs (dexamethasone, 0.2 mg/kg) to minimize brain swelling during the surgical procedure. The mouse scalp was shaved in the surgical region, and the mouse was placed into a stereotaxic frame with a heating pad. Lidocain was locally applied prior to the removal of the skin and periosteum above the skull. A 3 mm diameter craniotomy was performed above the right or left hemisphere using a dental drill (anteroposterior −2.2 mm; mediolateral +1.8 mm). The dura was carefully removed using fine forceps before aspiring the somatosensory cortex above the hippocampus using a vacuum pump connect to 29G blunt needle. The overlying alveus was carefully peeled away to expose the surface of the hippocampus. A custom-made metal tube sealed with a coverslip (#1) on the bottom side (both 3 mm in diameter) was inserted into the craniotomy and tightly fixed to the skull with acrylic glue. Once in place, the hippocampal window was fixed using ultraviolet light curable dental cement.

### *In vivo* imaging

After the surgery, mice received analgesics for 2 days (buprenorphine, 0.05 mg/kg, intraperitoneal injection) and be allowed to recover for at least 4 weeks before starting imaging sessions. During these sessions, mice were anesthetized under 4% isoflurane prior to be transferred to a custom-made 3D printed tiltable frame, based on ear bars and nose fixations, incorporating a mask delivering 1.5-2% isoflurane at 0.2 L/min O_2_. The eyes were protected with ointment (Bepanthen) and body temperature was maintained using a heating pad with anal probe. Imaging of CA1 pyramidal neurons was performed at 10–30 *μm* depth to avoid scare tissue at the surface while limiting optical aberrations steaming from the sample (19, 26). Typical image size was 20×20 *μ*m^2^ in *XY* with a pixel size of 20 nm, z stacks typically extended over 4 *μm* with a z-step size of 100 nm. Images were acquired with a 20 *μs* pixel dwell time, while excitation and STED laser powers were in the range of 10-20 mW after the objective lens. With these acquisition settings, no signs of phototoxicity were visible.

### PSF computation

The PSF of the STED beam was calculated using vectorial diffraction theory by Richard and Wolf (30, 31), which makes it possible to calculate the electromagnetic field (**E**) in an arbitrary point P close to the focal region of a high numerical aperture (NA≥ 0.7) objective, based on the Debye integral (32, 33):

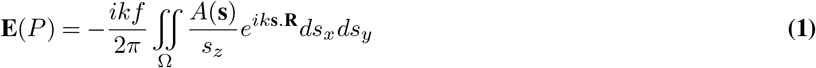

where k is the wavenumber, f the objective focal length, Ω the solid angle of the exit pupil from the focal spot, **s** the unit vector along each ray from the objective pupil to the focal volume, *A*(**s**) the complex amplitude of the incident laser beam after the objective and **R** the position vector of point P(*x, y, z*).

Considering the geometry depicted in Fig. 1a the diffraction integral can be expressed, in spherical coordinates, as (33, 34):

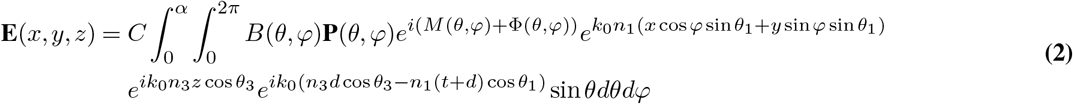

where C is a constant, *k*_0_ the wavenumber in the vacuum, *α* = arcsin(NA/*n*) the marginal ray angle, *n_l_* and *θ_l_* the refractive indices and incident angle in the immersion media (1), the coverslip (2) and the sample (3) respectively, *d* the depth in the sample, *t* the thickness of the coverslip, *B*(*θ,φ*) the amplitude profile of the incident beam, **P** the polarization state of the electromagnetic field in the focal region, *M* (*ρ, φ*) the phase profile of the input beam, corresponding in our case to the phase mask used to shape the STED beam, and *Φ*(*θ,φ*) is the wavefront distortion with respect to the Gaussian reference sphere, which describes the optical aberrations in the system. Note that, the second and third exponential terms correspond to the aberrations induced along the optical path through the coverslip and the biological sample (34).

**Fig. 1.**
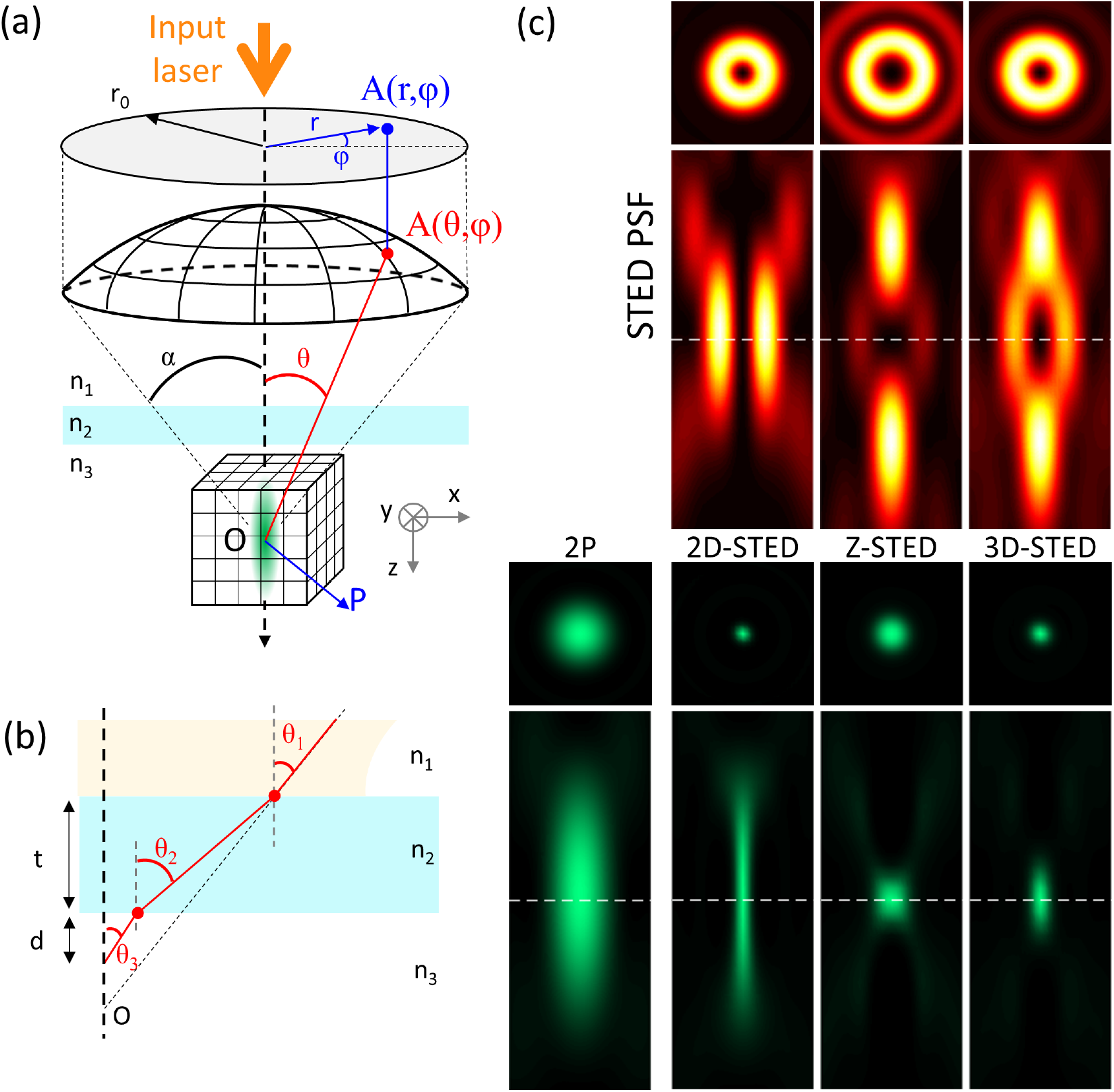
(a) Schematic representation of the propagation of a light wave focused by a high NA objective used to calculate the PSF in the vicinity of the focus. (b) Refraction angles within the coverslip. Due to the refractive index mismatch, each interface decreases transmission and induces spherical aberrations. (c) STED beam (top, fire LUT) and effective fluorescence (bottom, green LUT) PSFs in *XY* plane (square panels, image size 1×1 *μ*m^2^) and *XZ* plane (rectangle panels, image size 1×2.5 *μ*m^2^), for the different configurations used in the work (2-photon only, 2D-STED, z-STED and 3D-STED). The dashed line indicates the focus position.

While passing through an aplanatic objective, the incident plane wave transforms into a spherical wave converging to the focal point. Therefore, assuming a Gaussian profile of the input beam, the amplitude distribution after the objective can be expressed as:

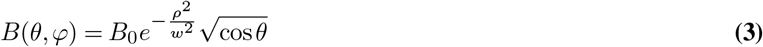

where *B*_0_ is a constant, *w* the beam waist, *ρ = f* sin *θ* the cylindrical coordinate on the exit pupil of the objective lens and 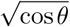 is the apodization term ensuring energy conservation while the beam pass through the objective. In addition, the objective transforms the input left-handed circular polarization **P**_0_(*θ,φ*) = (1,i, 0), classically used in STED microscopy, through tight focalization, which can be described as:

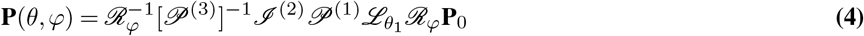

where 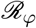 is the rotation matrix around z axis, 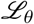 describe the change in electric field as it passes through the objective, 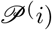 corresponds to the coordinate system rotation in medium *l* and 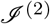 is the matrix describing the effect of the coverslip medium, considered as a stratified medium of two interfaces.

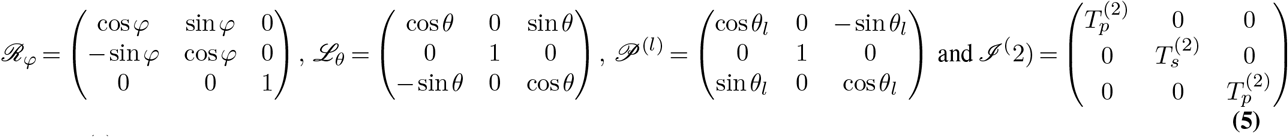

where 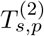 are the transmission coefficient in the coverslip (see (34) for complete derivation). Finally, in the case of left-handed circular polarization:

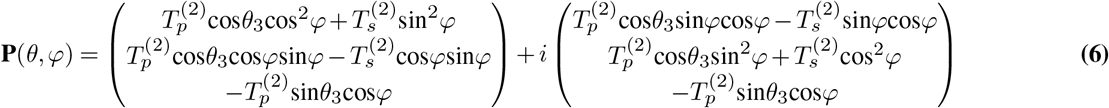

In the context of STED microscopy, the PSF of the STED beam is spatially shaped (Fig. 1c - top profiles) using specific phase masks *M*(*θ,φ*), that can be expressed as:

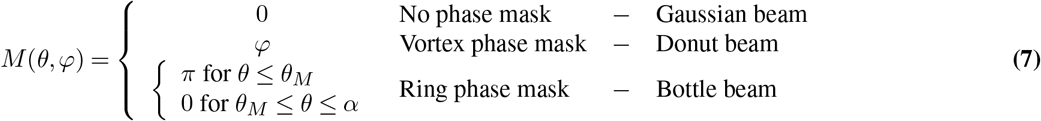

where 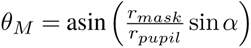 is the angle between the optical axis and the ray passing through the edge of the *π*-phase ring of the phase mask of radius *r_mask_* on the objective output pupil of radius *r_pupil_* (Fig. 2d - top panel).

**Fig. 2.**
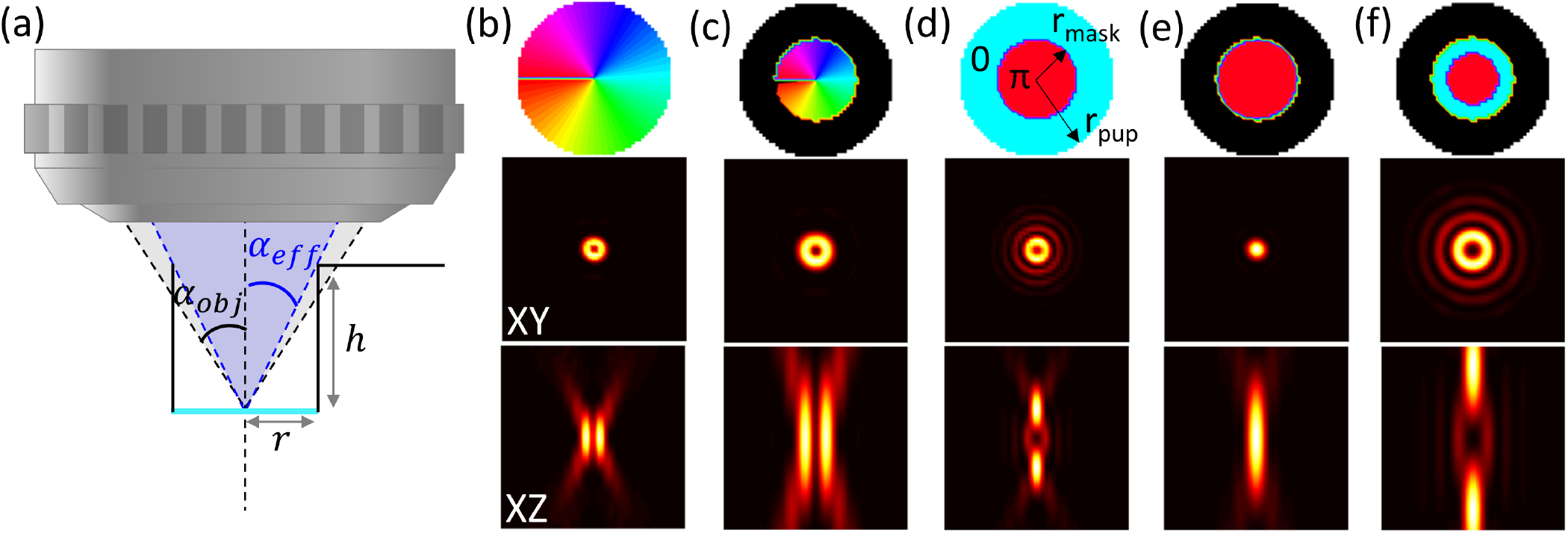
(a) Schematic of the hippocampal cranial window and its effect on the focused beam, notably the clipping of the outer optic rays. Numerical simulation of the STED beam *XY* (middle panels) and *XZ* (bottom panels) profiles calculated for the different phase masks (top panels) used in STED microscopy: Donut (b), effective donut in presence of hippocampal window (c), bottle beam (d), effective bottle beam in presence of hippocampal window (e) and same with adjusted phase mask radius (*r_mask_* = 0.33r_*pup*_) enabling the retrieval of the bottle profile (f). Image size: 5×5 *μ*m^2^.

The focal intensity can be calculated as the squared modulus of the electric field:

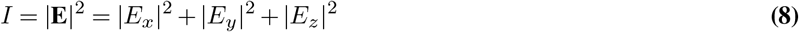

Lastly, the effective PSF (Fig. 1c - bottom profiles), is calculated as (35):

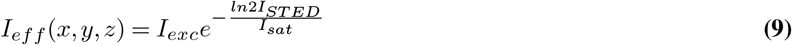

where *I_exc_* and *I_STED_* are the excitation and STED beams, respectively, and *I_sat_* is the saturation intensity, which describes the de-excitation rate of the molecules by the STED beam.

## Results and discussion

### Impact of the cranial window

The chronic hippocampal window used here was originally developed to image pyramidal neurons in the hippocampus by 2-photon microscopy (14, 15, 29) using a 0.8 NA objective. In this case, the specific geometry of the window, a metal cylinder sealed with a coverslip, limited the angle of the marginal rays transmitted through the window (Fig. 2.a). Indeed, to reach the hippocampus surface, the implanted cylinder and holder has a height (h) of 2.23 mm and an inner diameter (r) of 2.6 mm, which corresponds to a maximum opening angle of 30.2° and hence an effective NA of 0.67. While such an NA can be acceptable for 2-photon imaging, albeit at the expense of spatial resolution (notably axial resolution because of extended excitation PSF), it is prohibitive for STED microscopy. Indeed, beyond the NA limitation, the elimination of the marginal rays by the window design has a dramatic effect on the STED-PSF, rendering it useless, even counterproductive, for improving the STED axial resolution.

To further investigate this effect on the STED PSF and the resulting effective fluorescence PSF, we modified the calculation to consider the effect of the cranial window. We introduced an additional amplitude mask *T*(*ρ,φ*) in Eq. 2, which models the additional aperture stop at the entrance of the window leading to the clipping of the outer rays after the objectives. The diffraction integral can be expressed as:

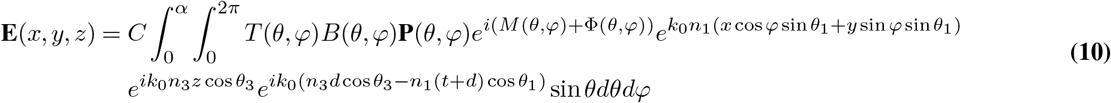

with

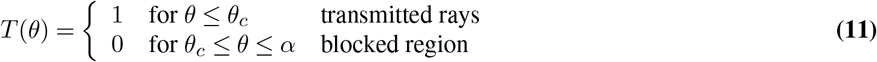

where *θ_c_* is the angle between the optical axis and the marginal ray of the cranial window on the output pupil of the objective. Fig. 2. (b-f) displays the results of these simulations. While the effect of the reduced NA is clearly visible on the donut PSFs (b and c), the window does not change the geometry of the phase mask and hence the shape of the PSF. Therefore, even if sub-optimal, this configuration still permits super-resolution imaging. In contrast, the bottle beam (d) is dramatically degraded in the presence of the cranial window: in the case of the ring phase mask, the outer rays (with 0 phase) are not passing through the hippocampal window, while the inner rays (with *π* phase) remain unaffected. This prevents destructive interference to happen in the focus and hence the formation of the central zero that is required for constricting the fluorescence around the edge of the fluorescence PSF, but leaving its central region free of STED light intensity (e). Note that, by simply adjusting the ring radius on the phase mask, one could effectively retrieve a correct bottle beam profile (f), yet with an elongated shape due to the limited effective NA, hence decreasing the axial resolution.

### New cranial window design and experimental validation

In order to retrieve a usable bottle beam and to achieve a subtantial STED gain in spatial resolution in all three dimensions, we designed a new cranial window, or hippocampal porthole. While increasing the radius of the conical window is possible (Fig. 2a), retrieving the full NA would require implanting a cylinder with a diameter of 5.1 mm into in the mouse brain, which is prohibitive in terms of the amount of cortical volume that would have to be surgically removed (about 45 mm^3^ of cortex). Instead, we chose to modify the geometry of the window.

Using a conical shape (Fig. 3a) makes it possible to benefit from the full nominal NA of the objective, while minimizing the amount of tissue that needs to be resected in order to expose the hippocampus, reducing it to about 14mm^3^, which is less than one third.

**Fig. 3.**
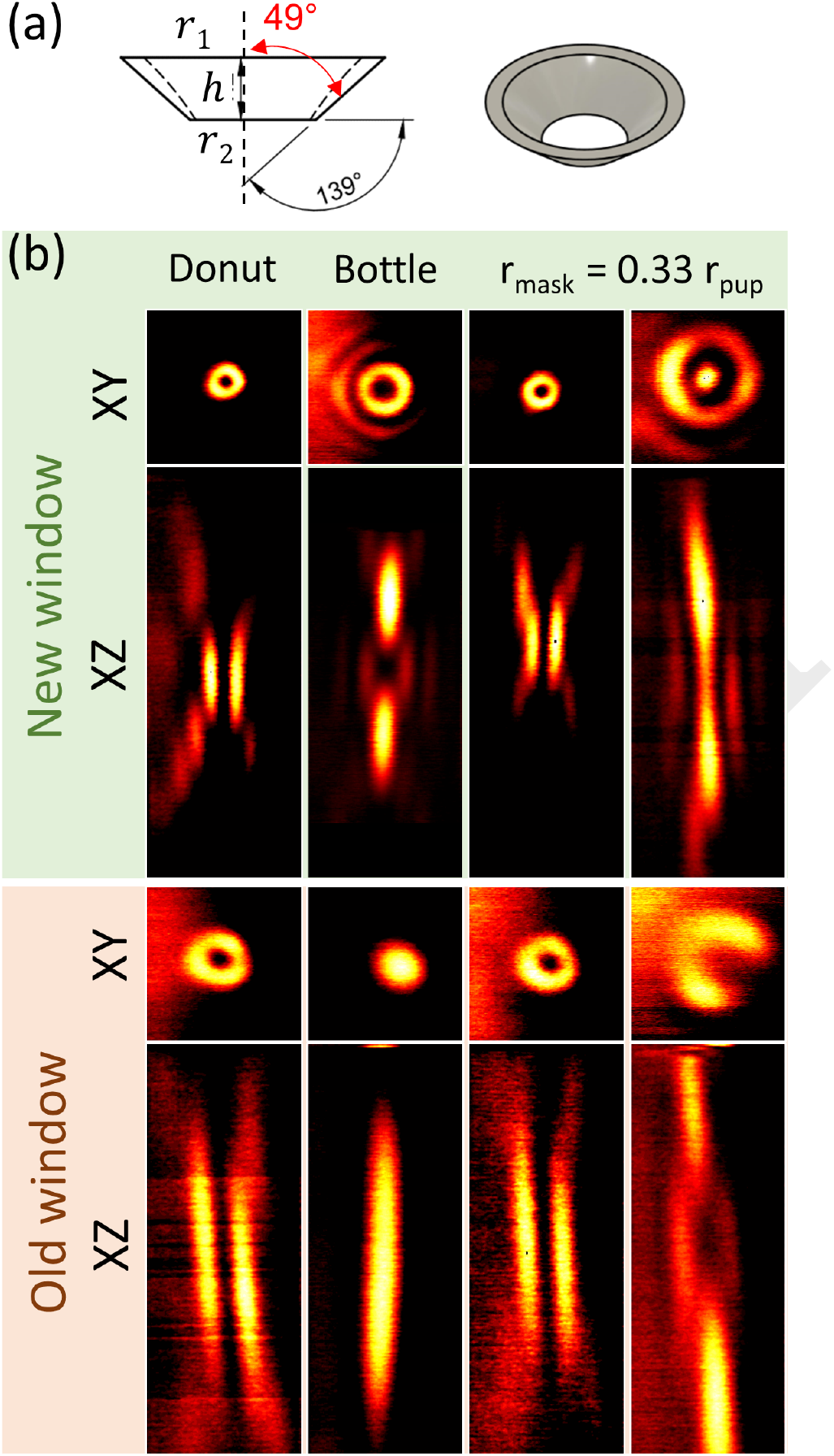
(a) Schematic of the new hippocampal window. The conical shape makes it possible to use the full NA of the objective while minimizing the cortical volume that needs to be removed. (b) STED beam PSFs in *XY* (2×2 *μ*m^2^) and *XZ* (2×8 *μ*m^2^) planes experimentally measured, using gold nano-particles attached to the coverslip, and imaged through the new (top panel) and old (bottom panel) cranial window designs, demonstrating the recovery of the appropriate bottle beam shape.

We first validated this cranial window geometry *ex vivo*, by placing gold nanoparticles on a poly-L-lysine coated coverslip, the same that we had used in the cranial window for *in vivo* imaging, and imaged them either through the cylinder (old window) or conical (new window) porthole without implantation on the head of the animal. This illustrates experimentally the impact of the cranial window design on the PSF of the STED beam.

Fig. 3b clearly illustrates the effect of the two different cranial window designs on the STED PSF. Beyond the reduction of the NA, the old cylindrical window seriously degrades the shape of the bottle beam, obliterating the central intensity minimum, which is a must for STED microscopy. By contrast, the new conical cranial window design permits the generation of optimal donut and bottle beam shapes. With the new design, the NA is limited by the optical design of the objective, and not the geometry of the optical access.

### *In vivo* 3D-STED imaging in the hippocampus

To visualize the gain in resolution in live conditions, fluorescence beads (*ø* 170 *μ*m) were attached to the coverslip using poly-L-lysine prior to be grafted into the animal. Imaging the fluorescent beads through the old and new window designs makes it possible to quantify and compare the gain in resolution between them. Fig. 4 displays images in *XY* and *XZ* direction of the fluorescent beads visualized through the cranial window. Note that the look-up table is adjusted between images for better visualization.

**Fig. 4.**
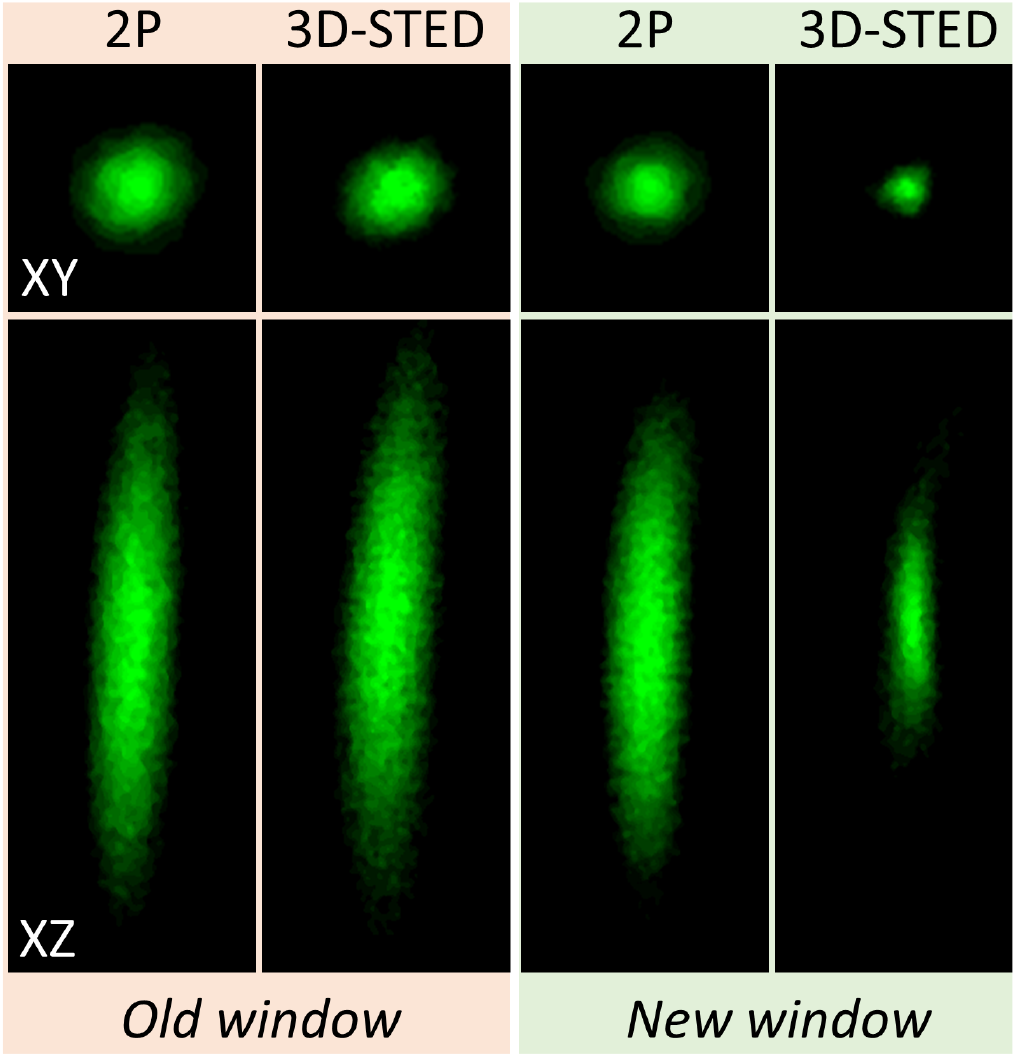
Effective fluorescence PSF in 2-photon and 3D-STED obtained by imaging the fluorescent nanoparticles attached below the coverslip in the old cylindrical and new conical cranial window implanted above the hippocampus of an adult mice. *XY* image size: 1.5×1.5 *μ*m^2^ and *XZ* image size 1.5×4 *μ*m^2^

Table 1 reports the axial and lateral resolution obtained experimentally using the two different windows together with the theoretical resolution obtained from numerical simulations. In this table, the spatial resolution is estimated as the full-width at half maximum (FWHM) of the PSF. Beyond the resolution, we quantified the maximum number of counts on the detector as a measure of image brightness (and thus SNR), which is strongly affected by the window geometry. Comparing our simulations with the experimental results, we normalized the number of counts in the simulated image with the value obtained from 2-photon images acquired with the new cranial window (which is expected to yield the highest number of counts).

**Table 1.** Spatial resolution (estimated as FWHM of the PSF) and maximum number of counts obtained in 2-photon and STED with the two different cranial window geometries. Mean ± SD from 10 fluorescent beads in 2 different samples prepared from the same batch.

|  |  | Lateral resolution (nm) |  | Axial resolution (nm) |  | Maximum intensity (# counts) |  |
| --- | --- | --- | --- | --- | --- | --- | --- |
|  |  | 2P | STED | 2P | STED | 2P | STED |
| Old window | Simulated | 472 | 456 | 2831 | 2788 | 60 | 17 |
| | Experimental | $470 \pm 10$ | $430 \pm 20$ | $2800 \pm 150$ | $2500 \pm 500$ | $50 \pm 20$ | $20 \pm 15$ |
| New window | Simulated | 344 | 81 | 1244 | 285 | 158 | 138 |
| | Experimental | $350 \pm 8$ | $80 \pm 10$ | $1200 \pm 100$ | $310 \pm 40$ | $158 \pm 8$ | $110 \pm 10$ |

The 2-photon PSFs are slightly improved by the new window, yielding a modest but clear improvement in spatial resolution. In contrast, the 3D-STED performance is greatly affected by the type of window design. With the old cylindrical window, the effective PSF is very similar to the 2-photon PSF but with a strongly reduced signal, as expected from the absence of a zero in the bottle beam profile, diminishing the excitation of molecules without yielding of a gain in spatial resolution. In contrast, the new conical window permits a significant constriction of the fluorescent spot, while largely preserving the signal level.

Having established this proof of principle, we validated this modified hippocampal window on biological samples by imaging fluorescently labelled neurons in living transgenic mice. Fig. 5a shows a segment of dendrite in the *stratum radiatum* of the CA1 region in the hippocampus of a living mouse. Notably, the fine morphological features, including the hallmark cup-like shapes of spine heads and the ultrathin neck regions, connecting the spine head with the dendrite can be appreciated with unprecedentedly high image quality in an *in vivo* setting. Fig. 5b and c show a volume rendering of the same segment of dendrite qualitatively illustrating the gain in resolution and anatomical fidelity that can be achieved by our improved approach. Finally, we also quantified neck diameters of the dendritic spines using a semi-automatic software (36), which was specifically designed for morphometric analysis of super-resolution images of dendritic spines. The results are summarized in Table 2 and are consistent with the published literature based on electron microscopy or STED imaging in brain slices (37–39).

**Fig. 5.**
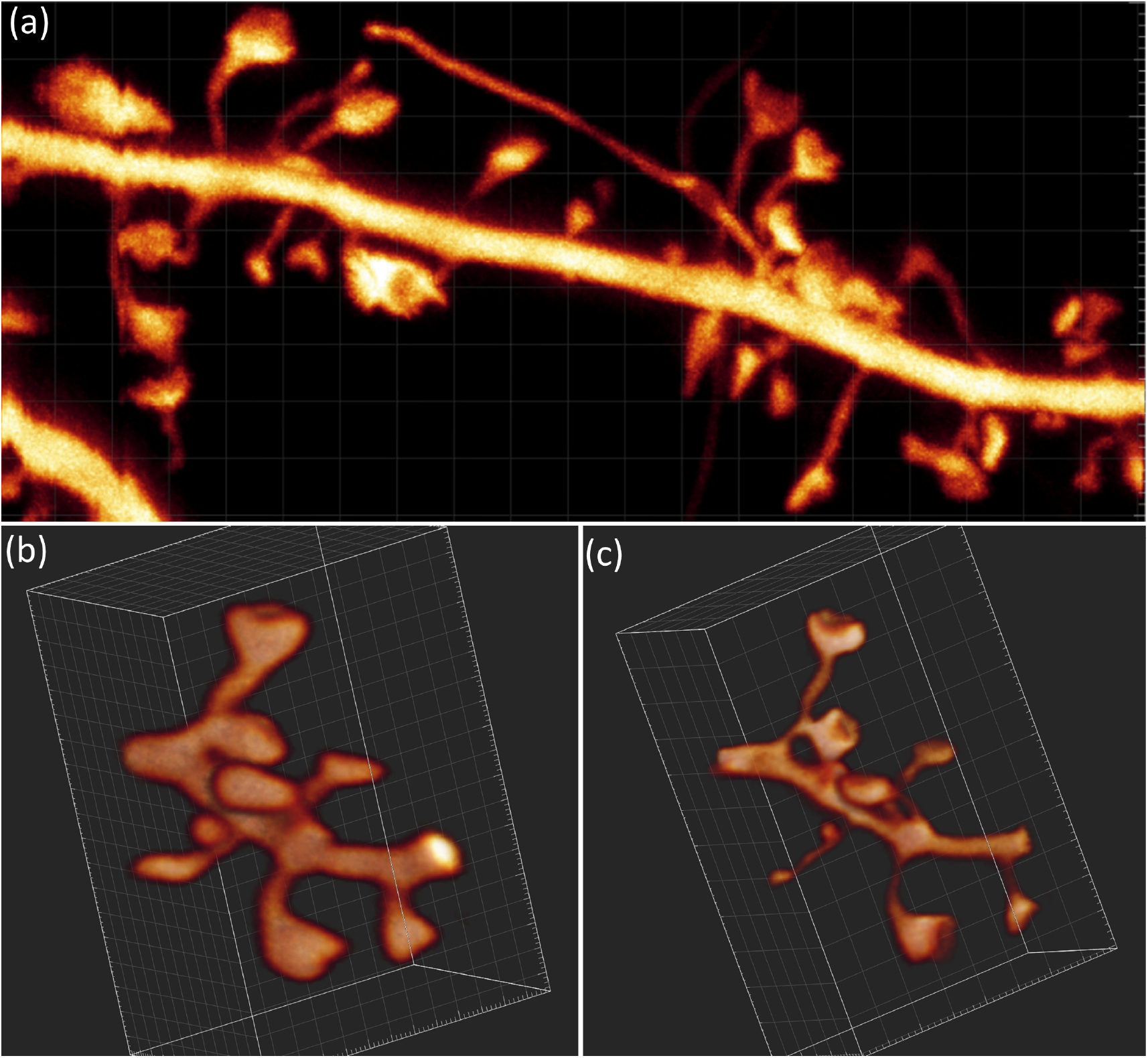
(a) Image of a YFP labelled segment dendrite in the *stratum radiatum* of a *Thy*1–*H^tg/+^*mouse, lying about 30 *μm* below the surgically created surface. Image size 5×11 *μ*m^2^. (b) and (c) 3D rendering of the same segment of dendrite obtained with 2-photon and 3D-STED imaging, respectively. Image size 5×3.2×3.2 *μ*m^3^.

**Table 2.**
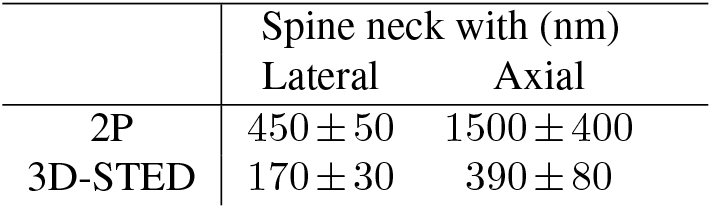
Average of thinnest spine neck widths, measured in the lateral and axial directions by 2-photon and 3D-STED microscopy. Mean ± SD from 23 dendritic spines collected from 3 mice.

## Conclusion

In this work, we propose and validate a modified hippocampal window design, which makes full use of the NA of a long-working distance objective. While this new design by itself already increases the spatial resolution and optical sectioning of 2-photon microscopy, it is essential for STED microscopy. Notably, it can preserve the bottle beam shape needed for 3D-STED microscopy. We illustrate the benefit of this new cranial window design by visualizing dendritic spines, greatly improving STED image quality, rendering it comparable to the state of the art in brain slice preparations.

Our new approach improves the achievable spatial resolution for nanoscale imaging of neuroanatomical structures and compartments (e.g., the extracellular space between brain cells), enabling longitudinal investigations into how their dynamics may underpin the ability of neurons and their networks to adapt themselves to ever-changing environmental conditions in health and disease.

## ACKNOWLEDGEMENTS

This project received funding from the European Union’s Horizon 2020 research and innovation program under the *Marie Sklodowska-Curie Grant*, Agreement No. 794492, as well as from the *Fonds AXA pour la Recherche, AXA Banque DIrection Banque Patrimoniale et ses donateurs* to SB, from *Doctoral School for Health and Life Sciences* (a PhD Fellowship to JR), the *European Research Council* (ERC-SyG ENSEMBLE # 951294), *Human Frontiers Science Program* (# RGP0036/2020), *ERA-NET NEURON* (ANR-17-NEU3-0005) and *Agence Nationale de la Recherche* (ANR-17-CE37-0011) to UVN. We thank the animal facility at IINS for their support.

## Notes

### Competing Interest Statement

The authors have declared no competing interest.

